# Cell-fate plasticity, adhesion and cell sorting complementarily establish a sharp midbrain-hindbrain boundary

**DOI:** 10.1101/857870

**Authors:** Gokul Kesavan, Stefan Hans, Michael Brand

## Abstract

The formation and maintenance of sharp boundaries between groups of cells play a vital role during embryonic development as they serve to compartmentalize cells with similar fates. Some of these boundaries also act as organizers, with the ability to induce specific cell fates and morphogenesis in the surrounding cells. The midbrain-hindbrain boundary (MHB) is an example of such an organizer that also acts as a lineage restriction boundary that prevents the intermingling of cells with different developmental fates. However, the mechanisms underlying the lineage restriction process remain unclear. Here, using a combination of novel fluorescent knock-in reporters, live imaging, Cre/lox-mediated lineage tracing, atomic force microscopy-based cell adhesion assays, and mutant analysis, we analyze the process of lineage restriction at the MHB and provide mechanistic details. Specifically, we show that lineage restriction occurs by the end of gastrulation, and that the subsequent formation of sharp gene expression boundaries in the developing MHB occur through complementary mechanisms, namely cell-fate plasticity and cell sorting. Further, we show that cell sorting at the MHB involves differential adhesion among midbrain and hindbrain cells that is mediated by N-cadherin and Eph-Ephrin signaling.

## Introduction

The concept of boundaries between gene expression domains is central and crucial to our current understanding of organ development because some of these boundaries also act as organizers or local signaling centers (Kiecker and Lumsden, 2005; Dahmann et al., 2011). Organizers are groups of cells that instruct other cells in their vicinity to acquire specific developmental fates and this fundamental process is essential for proper embryonic development (Arias and Steventon, 2018). The midbrain-hindbrain boundary (MHB), also known as the isthmic organizer (midbrain-hindbrain organizer) is an example of such an organizer that forms at the interface of the midbrain (mesencephalon, mes) and the hindbrain (cerebellum, metencephalon, met), and is vital for the formation and function of both the midbrain and the cerebellum (Rhinn et al., 2006; Gibbs et al., 2017). Cells at these signaling centers secrete various morphogens like Wnt and fibroblast growth factors (Fgf) which provide “positional information” and help establish proper tissue patterning and cell fate commitment (Gibbs et al., 2017).

In vertebrates, the interface between the expression domains of the two transcription factors, namely, Otx and Gbx, constitutes the MHB, and in zebrafish, the MHB develops from the initially-overlapping expression domains of Otx2 and Gbx1 that subsequently sort out and form a sharp boundary (Raible and Brand, 2004; Rhinn et al., 2003). During this process, morphogens such as Wnt, Fgf, and transcription factors, like Engrailed1/2, and Pax2/5/8, sequentially induce MHB formation, and their subsequent interplay is critical for the maintenance of the MHB (Rhinn and Brand, 2001; Wurst and Bally-Cuif, 2001; Raible and Brand, 2004; Rhinn et al., 2006; Dworkin and Jane, 2013). The above-mentioned factors (Otx, Gbx, Wnt1, Fgf8, Pax, and Eng) comprise the core of the MHB signaling machinery and any functional disruption of these factors interferes with patterning at the MHB (Dworkin and Jane, 2013; Gibbs et al., 2017).

Several studies using various vertebrate models have addressed whether the MHB also acts as a lineage restriction boundary, apart from its role as an organizer, and most studies argue in favor of the MHB also being a lineage restriction boundary (Zervas et al., 2004, Sunmonu et al., 2011). We have previously demonstrated the existence of a lineage restriction boundary at the zebrafish MHB using a combination of high-resolution time-lapse imaging, single-cell labelling, and transplantation experiments (Langenberg and Brand, 2005; Langenberg et al., 2006). Our observations imply a cell-cell communication mechanism that restricts the migration of cells across the presumptive MHB but still permits cell movement within the group of cells on either side of the boundary. Thus, while it is known that Otx2 and Gbx1 establish their expression domains on either side of the MHB, the mechanisms by which this Otx-Gbx interface acts as a lineage restriction boundary, and the cell biological processes that prevent the intermingling of cells destined to dissimilar developmental fates, remain poorly understood.

Therefore, we have addressed these questions using the developing zebrafish as a model. Specially, we visualized the establishment of the MHB in real-time by generating various novel fluorescent reporter and Cre driver lines for *otx2b*, *wnt1*, *gbx1*, *gbx2*, and *fgf8a* using CRISPR/Cas9-mediated knock-in strategies (Kesavan et al., 2017; Kesavan et al., 2018). Using a combination of time-lapse imaging and Cre/lox-mediated lineage tracing, we show that lineage restriction at the MHB occurs by the end of gastrulation. Next, we demonstrate that cells indeed sort at the MHB to establish a sharp boundary and that the underlying molecular mechanisms include differential adhesion between prospective midbrain and hindbrain progenitors that involve N-cadherin and signaling through the Eph-ephrin pathway.

## Results

### Initially overlapping gene expression domains segregate overtime

The gene expression boundary abutting the *otx2b*-*gbx1* expression domains demarcates the MHB primordium. In order to visualize these developmental events in real time, we generated CRISPR/Cas9-based knock-in zebrafish lines. The coding sequences for the reporters were knocked-in upstream of the corresponding ATG and, thus, under the control of the endogenous promoter/enhancer elements (Fig 1A, schematic representation). Importantly, as shown previously, reporter expression faithfully reproduces the endogenous gene expression patterns (Kesavan et al., 2017; Kesavan et al., 2018).

**Figure 1.**
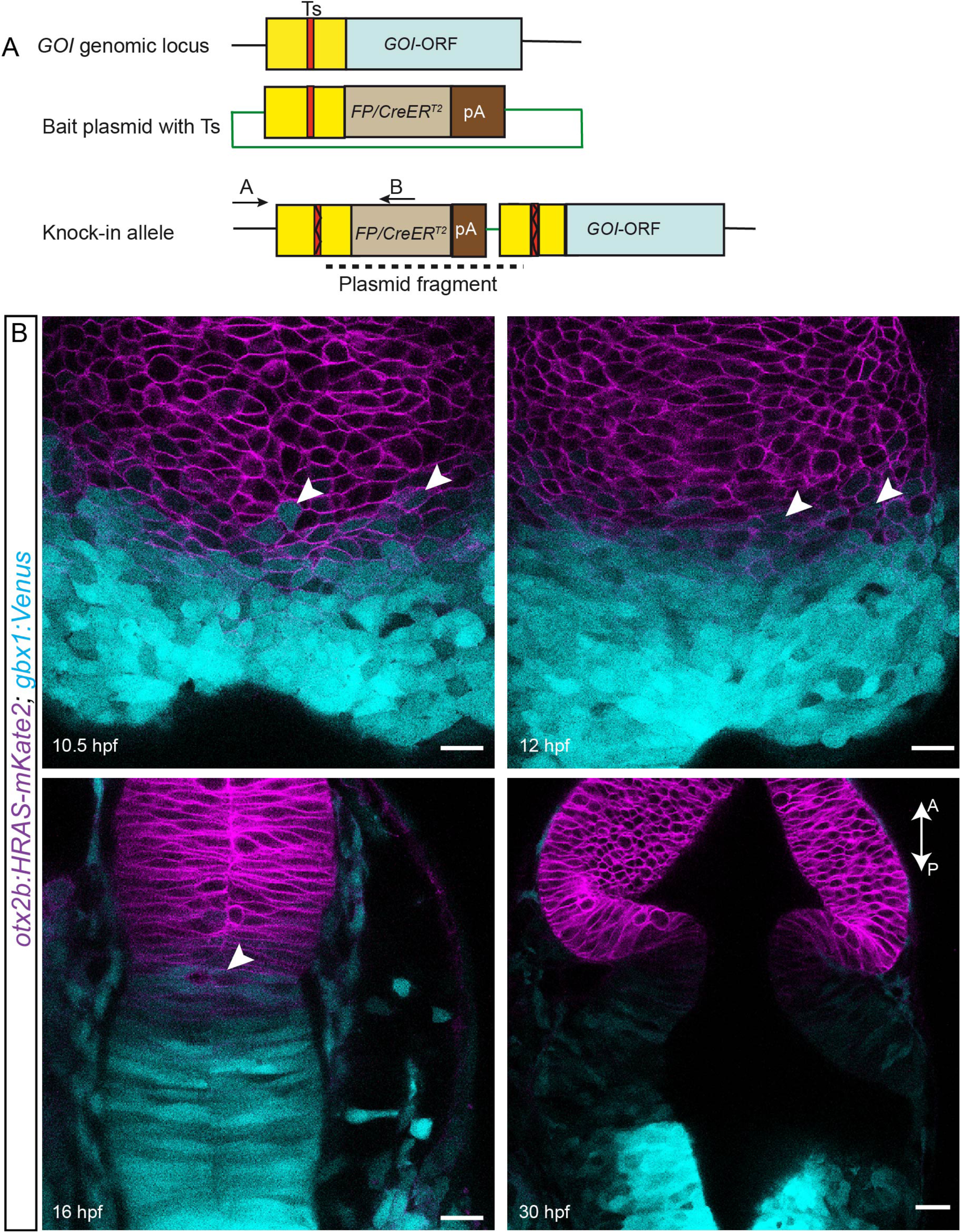
Initially overlapping expression domains segregate overtime. (A) The knock-in strategy used for generating transgenic fish is schematized. A target site (Ts) located upstream of the open reading frame (ORF) of a gene of interest (GOI) is chosen. A bait plasmid is constructed by cloning the sequence upstream to the ORF including the target site. The bait plasmid, *sgRNA* against the target site, and *Cas9* mRNA are injected into the embryo at the 1-cell stage. The Cas9 protein creates double strand breaks at both Ts in the genomic locus and in the bait, plasmid followed by integration of the linearized bait plasmid. Only integration in the forward orientation will result in fluorescent reporter or *CreER_T2_* expression. The primer pair (A+B) can be used to screen for and verify precise integration of the plasmid. The forward primer A is located outside of the bait and the reverse primer B is located within the fluorescent reporter/*CreER_T2_* sequence. (B) Live imaging was used to follow the expression of both mKate2 (membrane localized) and Venus, driven by the *otx2b* and *gbx1* locus respectively, at the various time points indicated. Overlapping expression boundaries were initially observed during the segmentation stages, i.e., between 10.5 to 16 hpf, and these segregated over time, based on changes in the presence of double positive cells (mKate2 and Venus positive). Scale bar 20 µm (B). GOI, Gene of interest; FP, fluorescent protein, the anterior-posterior axis of the embryos are marked with A-P and an arrow.

To address Otx-Gbx boundary formation in real time, we used time-lapse imaging of *otx2b:HRAS-mKate2*; *gbx1:Venus* double transgenic embryos labelling cell membranes in the midbrain and cells in the hindbrain, respectively. However, this process could only be visualized from tail bud stage onwards (10 hours post fertilization, hpf) because this delay corresponds to the time required for the reporter (mKate2 or Venus) to become functional, despite being fast-folding proteins. Nonetheless, an overlap in the expression of *otx2b* (midbrain, mKate2) and *gbx1* (hindbrain, Venus) can be observed at the neural plate at the 1-2 somite stage (10.5-11 hpf) as evidenced by the presence of cells that are double positive for both mKate2 and Venus (Fig 1B). Importantly, these cells could only be observed close to the prospective MHB and were not present in the deeper cell layers of the expression domains of either *otx2b* or *gbx1* (Fig 1B). Subsequently, around the 10-14 somite stage (15-16 hpf), this intermingling of double-positive midbrain hindbrain cells resolved into a sharper boundary (Fig 1B, supplementary movie 1). Overlapping expression domains were completely absent by around 30 hpf. Additionally, at this time point, the double positive cells, along with gaps that represent *otx2b*-derived cells within hindbrain domain, were also not visible (Fig 1B, supplementary movie 1). Together, these observations imply a sequence of events during MHB formation wherein an initial overlap in the expression domains of Otx and Gbx resolves into a sharp, non-overlapping boundary over time. The process by which such initially overlapping gene expression domains segregate to form sharp boundaries can involve multiple mechanisms, such as loss of cell identity (cell-fate plasticity) or cell sorting (Batlle and Wilkinson, 2012). Hence, we next investigated if these mechanisms play a role in MHB formation.

### Lineage restriction at the MHB occurs at the end of gastrulation

To understand changes in cell identity, i.e., the establishment of lineage restriction at the midbrain hindbrain boundary (MHB), we attempted two types of lineage tracing, namely, short-term or transient labelling, and long-term or permanent tracing. We used both approaches in order to derive additional information on potential identity change or fate-plasticity in the cells abutting the MHB. First, for short-term tracing, CRISPR/Cas9-mediated knock-ins of fluorescent reporters for multiple genes expressed at the MHB, namely, *otx2b*, *wnt1*, *gbx1*, and *gbx2* were generated (Fig 2C, schematic representation of gene expression boundaries). Next, to follow the fate of Otx2 and Wnt1 expressing cells at the MHB, these fluorescent reporter lines were tagged with a nuclear localization signal (NLS), to generate *otx2b:Venus-NLS* and *wnt1:Venus-NLS* fishes. Venus was used as the reporter because it is fast folding, which allows visualization of the early events in MHB formation. Moreover, it has a known lifetime of about 24 hours (Li et al., 1998., Snapp, 2009), such that the ‘perdurance’ of Venus protein, even when the promoter is turned off, can be used as a lineage tracing marker.

**Figure 2.**
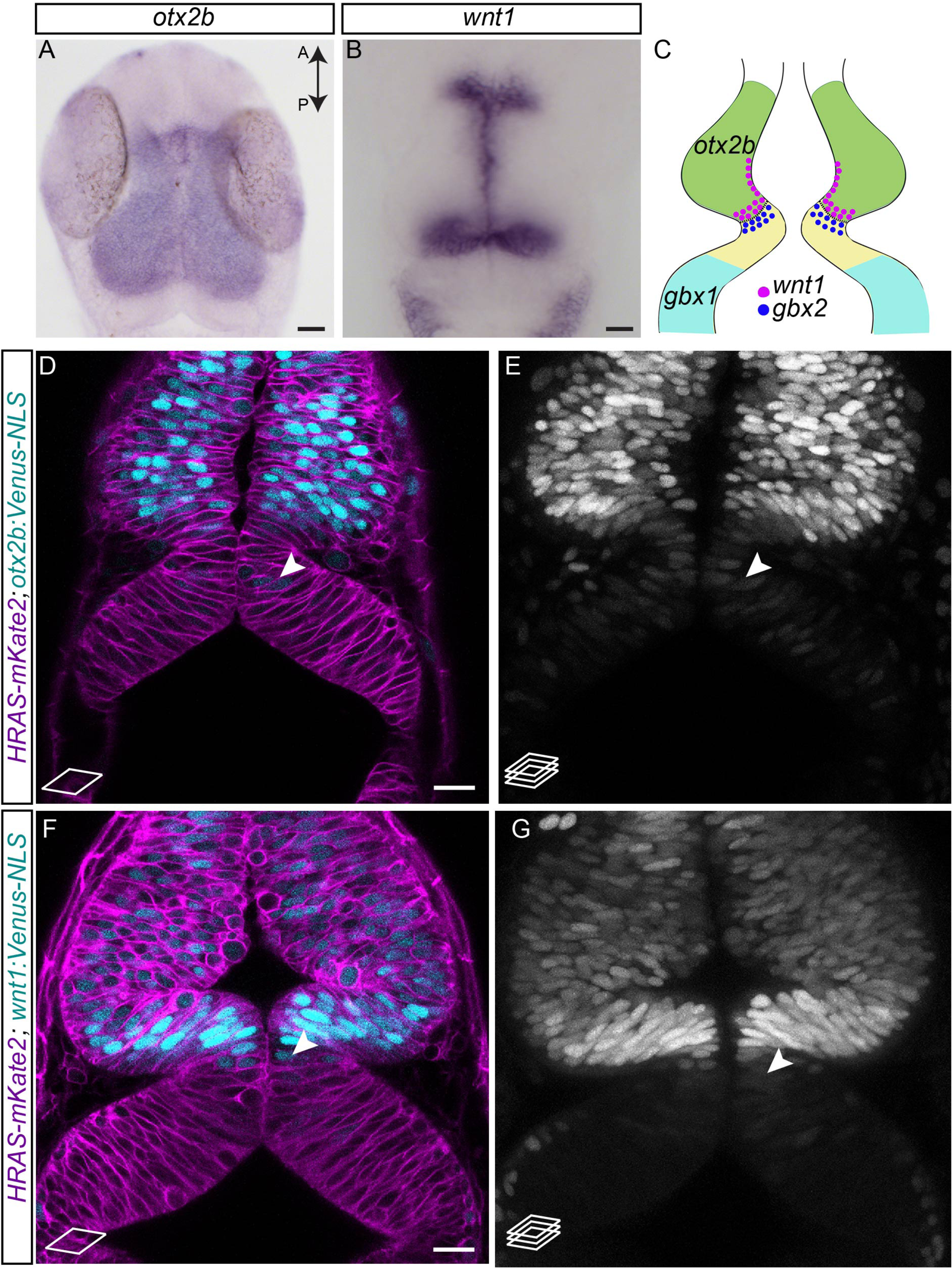
Lineage restriction in the midbrain occurs at the end of gastrulation. (A and B) Whole mount *in situ* hybridization (flat mount) in 24 hpf embryos for midbrain markers, namely *otx2b* and *wnt1*, show that their expression domains abut sharply at the MHB at this time point. (C) Schematic representation of the various marker genes expressed at the MHB in a 24 hpf embryo. Midbrain specific genes are *otx2b* (green) and *wnt1* (magenta dots), hindbrain specific genes are *gbx1* (cyan) and *gbx2* (blue dots). (D-G) Venus fluorescent protein expression at 24 hpf in live embryos. Membrane-localizing mkate2 mRNA (red fluorescent protein) was injected into 1-cell stage embryos to ubiquitously label all cells and visualize tissue architecture. (D-E) *otx2b:Venus*-positive cells were not only present in the midbrain domain but also in the hindbrain domain (arrowheads). (F-G) Similarly, *wnt1:Venus*-positive cells were present in the midbrain domain and were observed in the hindbrain domain (arrowheads). Images are from a single confocal plane in panels D and F, while the maximum projection images are shown in panels E and G. Scale bar 10 µm (A-B), 20 µm (D-G).

*in situ* hybridization at 24 hpf showed that *otx2b* and *wnt1* mRNA expression was restricted to the midbrain region with a sharp boundary abutting the MHB (Fig 2A and 2B). In contrast, at 24 hpf, in the *otx2b:Venus* and the *wnt1:Venus* reporter embryos, the presence of Venus fluorescent protein (perdurance) was observed in the hindbrain domain, albeit with a weaker intensity than in the midbrain domain (Fig 2D-2G), suggesting that these cells had indeed expressed *otx2b* and *wnt1* at an earlier time point during MHB development. Further, time-lapse imaging of *otx2b:Venus-NLS* and *wnt1:Venus-NLS* embryos not only showed the presence of Venus-positive cells in the hindbrain domain, but also that these cells remained within this domain over time (Supplementary movies 2 and 3). Taken together, these observations imply that cells abutting the MHB are indeed capable of changing their gene expression and may display cell-fate plasticity.

Next, to verify if a similar mechanism operates in hindbrain cells as well, hindbrain boundary markers like *gbx1* and *gbx2* were lineage traced using *gbx1:Venus* and *gbx2:Venus-NLS* reporter lines. *in situ* hybridization at 24 hpf for hindbrain markers, namely *gbx1* and *gbx2*, showed sharp expression boundaries posterior to rhombomere 1 for *gbx1* and at the MHB for *gbx2* (Fig 3A and 3B). In contrast, at 24 hpf in the reporter fish, the presence of Venus fluorescent protein was observed in the midbrain domain, albeit with relatively weaker fluorescence intensity than that seen in the hindbrain domain (Fig 3C-3F). Further, as shown in supplementary movie 1, these *gbx1*:*Venus*-positive cells were also *otx2b*-positive and remained in the midbrain domain. Thus, these observations point to a pattern of perdurance of hindbrain markers (*gbx1*) in the midbrain domain and of midbrain markers (*otx2b* and *wnt1*) in the hindbrain domain.

**Figure 3.**
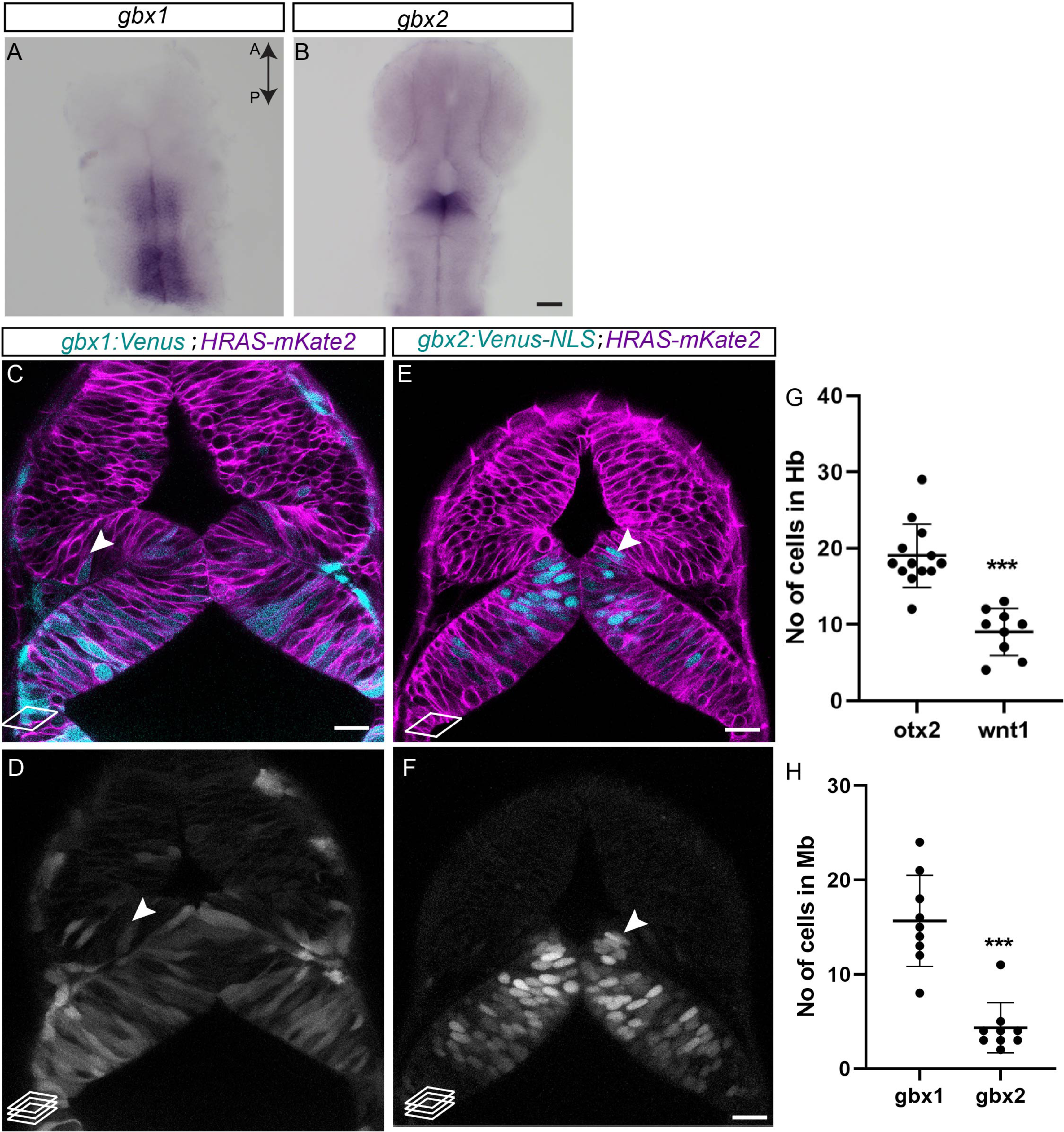
Lineage restriction in the hindbrain also occurs at the end of gastrulation. (A and B) Whole mount *in situ* hybridization (flat mount) in 24 hpf embryos for hindbrain markers, namely *gbx1* and *gbx2*, show sharp expression boundaries posterior to rhombomere 1 for *gbx1* and at the MHB for *gbx2* at this time point. (C-F) Venus fluorescent protein expression at 24 hpf in live embryos. Membrane-localizing mkate2 mRNA (a red fluorescent protein) was injected into 1-cell stage embryos to ubiquitously label all cells and visualize tissue architecture. (C-D) In contrast to expression patterns seen in *in situ hybridizations*, the *gbx1:Venus* transgenic line showed Venus-positive cells in the midbrain domain (arrowheads). (E-F) In the *gbx2:Venus* transgenic line, very few Venus-positive cells could be observed in the midbrain domain, especially at the caudal midbrain domain (arrowheads). Images are from a single confocal plane in panels C and E, and the maximum projection images are shown in panels D and F. (G-H) Quantification of Venus-positive cells in their non-expression domains, i.e., *otx2b* and *wnt1* in the hindbrain, and *gbx1*, *gbx2* in the midbrain, shows that greater numbers of *otx2* and *gbx1* cells were present in their non-expression domains, compared to *wnt1* and *gbx2*, respectively. The two-tailed, unpaired ‘*t*’-test was used to calculate statistical significance, and each point in the graph represents an individual embryo. *otx2* (n=13) vs *wnt1*(n=9), p< 0.0001, *gbx1* (n=9) vs *gbx2* (n=9), p<0.0001. 10 µm (A-B), 20 µm (C-F).

Next, we quantified this perdurance and show that, on average per embryo, 19 *otx2b*:Venus-NLS-derived cells and 9 *wnt1*-derived cells were found in the hindbrain domain (Fig 3G), i.e., greater numbers of *otx2b*-derived cells showed perdurance than *wnt1*-derived cells. Similarly, while there were 15 *gbx-1* derived cells in the midbrain, there were only 4 *gbx2*-derived cells (Fig 3H). To understand why there may be more *otx2b* than *wnt1* cells, or more *gbx1* than *gbx2* cells, we evaluated gene expression onset by whole mount *in situ* hybridization. The results revealed that *otx2b* expression occurs earlier than that of *wnt1* (6 hpf vs 10 hpf); likewise, *gbx1* expression occurs earlier (early onset) than that of *gbx2* (6 hpf vs. 10 hpf) (Supplementary Fig S1), indicating that genes with earlier onset may show greater plasticity compared to genes with later onset. Further, these observations, when combined with the pattern of perdurance seen above, imply that lineage restriction occurs in both midbrain and hindbrain cells at the end of gastrulation.

Next, to substantiate that lineage restriction may occur at the end of gastrulation, we permanently labelled cells in the MHB region by generating CRISPR/Cas9-based CreER_T2_ knock-in lines and crossing them with a zebrabow responder line (Pan et al., 2013). Zebrabow-based lineage tracing allows stochastic recombination, and the resultant fluorescent protein combinations provide an opportunity to understand lineage decisions and clonal origins with high temporal resolution. CreER_T2_-mediated recombination was induced with 4-hydroxy tamoxifen, administered at 6 hpf and/or 24 hpf, and all embryos were imaged at 48 hpf (Fig 4A, schematic representation). In the *otx2b:CreER_T2_* line combined with the zebrabow line, several recombined cells were observed in the hindbrain domain when 4-hydroxy tamoxifen was used at 6 hpf. In contrast, no recombined cells were seen after 4-hydroxy tamoxifen induction at 24 hpf (Fig 4B-G). Similarly, the *fgf8a:CreER_T2_* crossed with the zebrabow line revealed the presence of fgf8a-derived cells in the midbrain with 4-hydroxy tamoxifen induction at 6 hpf (Fig 4H-4J). 4-hydroxy tamoxifen induction at 24 hpf in this line did not yield a visible readout, probably because of low recombination efficiency and low Fgf8 expression. These permanent lineage tracing results not only concur with the perdurance data described above, but also suggest that (i) lineage restriction indeed occurs at the end of gastrulationand that (ii) there must be other mechanisms, such as cell sorting, that serve to subsequently establish and maintain sharp expression boundaries, i.e., from the tail bud stage onwards.

**Figure 4.**
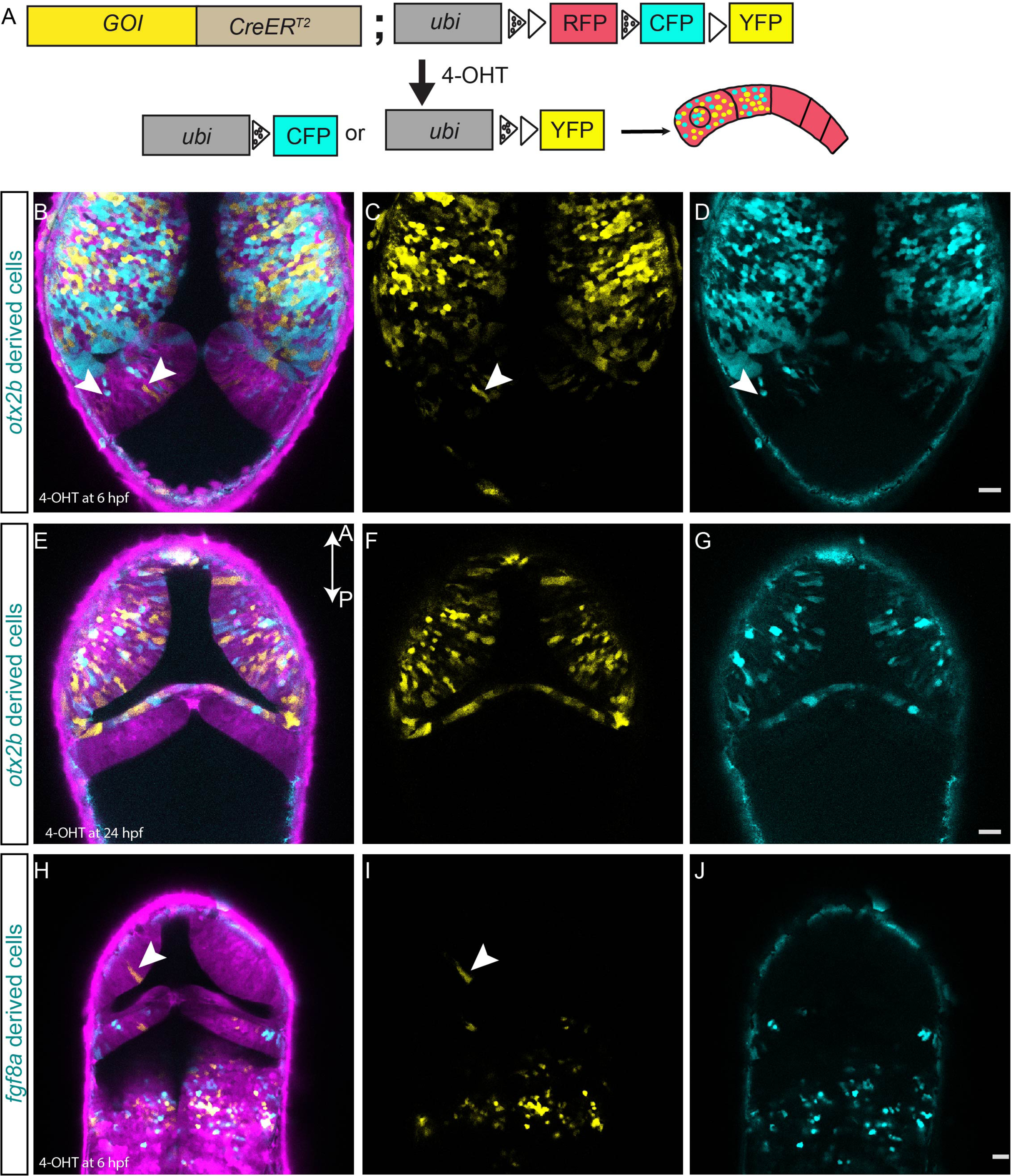
Zebrabow-based lineage tracing to visualize lineage restriction patterns at the MHB. Multicolor-labeling of midbrain and hindbrain cells using the *otx2b:CreER_T2_* and *fgf8a:CreER_T2_* knock-in driver lines. (A) Schematic representation of *CreER_T2_*-mediated recombination strategy using the zebrabow transgenic responder fish. In cells expressing CreER_T2_, 4-OH-tamoxifen (4-OHT) induces recombination between either the two *lox2272* sites (marked by dotted triangles) or the two *loxp* sites (marked by a triangle), which results in the stochastic labeling of cells due to the expression of CFP (cyan fluorescent protein) or YFP in the recombined cells. All non-recombined cells express only RFP (red fluorescent protein). In a cell with multiple copies of RFP, CFP, and YFP, CreER_T2_*-*mediated stochastic recombination events lead to the formation of clones marked by different colors. Embryos obtained by crossing Cre driver fish (either *otx2b* or *fgf8a*) with the zebrabow responder line were treated with 4-OH-tamoxifen, either at 6 hpf (1 μM) or 24 hpf (10 μM) for 12 h, and these embryos live-imaged at 48 hpf. (B-D) *otx2b:CreER*_T2_ embryos treated with Tam at 6 hpf show effective recombination in the midbrain region but also a few recombined cells in the hindbrain region (arrowheads). (E-G) *otx2b:CreER*_T2_ embryos treated with Tam at 24 hpf show recombined cells only in the midbrain. (H-J) Embryos of the *fgf8a:CreER _T2_* knock-in driver line treated with Tam at 6 hpf show effective recombination in the hindbrain with an exception of few recombined cells in the midbrain (arrowheads). Scale bar 20 µm (B-J).

### Cell sorting at the MHB

We looked at cell sorting as a possible mechanism of MHB formation and maintenance after lineage restriction was established at the tail bud stage. To track boundary cells with greater sensitivity in real time, we live imaged *wnt1:Venus*-*NLS* fluorescent reporter fish from the 1-2 somites stage till the 6-7 somites stage (10.5-12 hpf). We used Wnt1 as the marker because it accurately identifies midbrain boundary cells, and as mentioned earlier, imaging before 10 hpf was not possible due to the time required for reporter maturation. Using z-stacks of confocal images acquired over a narrow time interval (every 150 seconds), we were able to track individual cells (using labelled nuclei) with high spatial and temporal resolution. All cells migrated only anteriorly and individual cell tracking of the *wnt1* cells showed zig-zag movements and crossovers among these cells (Fig 5B and supplementary movie 5). Further, from around 14 hpf when the neural rod has formed, cell movement was restricted, and only inter-kinetic nuclear migration is observed (supplementary Movie 3), rather than active whole-cell movement. Interestingly, we could identify individual cells that were initially distant from the group of *wnt1*:*Venus-NLS*-positive boundary cells migrating anteriorly (Fig 5A, arrowhead). We tracked the movement of one such straggling *wnt1*:*Venus-NLS*-positive cell and found that this cell actively migrated towards the group of other boundary cells (Fig 5A, supplementary movie 4). Such active migration of straggling *wnt1* cells was observed in multiple embryos (in 5 out of 8 movies imaged; an additional time-lapse movie depicting such behavior is shown in supplementary movie 6). Thus, it appears that cell sorting during MHB formation involves active cell mixing and migration of cells both within and between brain compartments, both of which may contribute to establishing sharp gene expression boundaries during neural rod formation.

**Figure 5.**
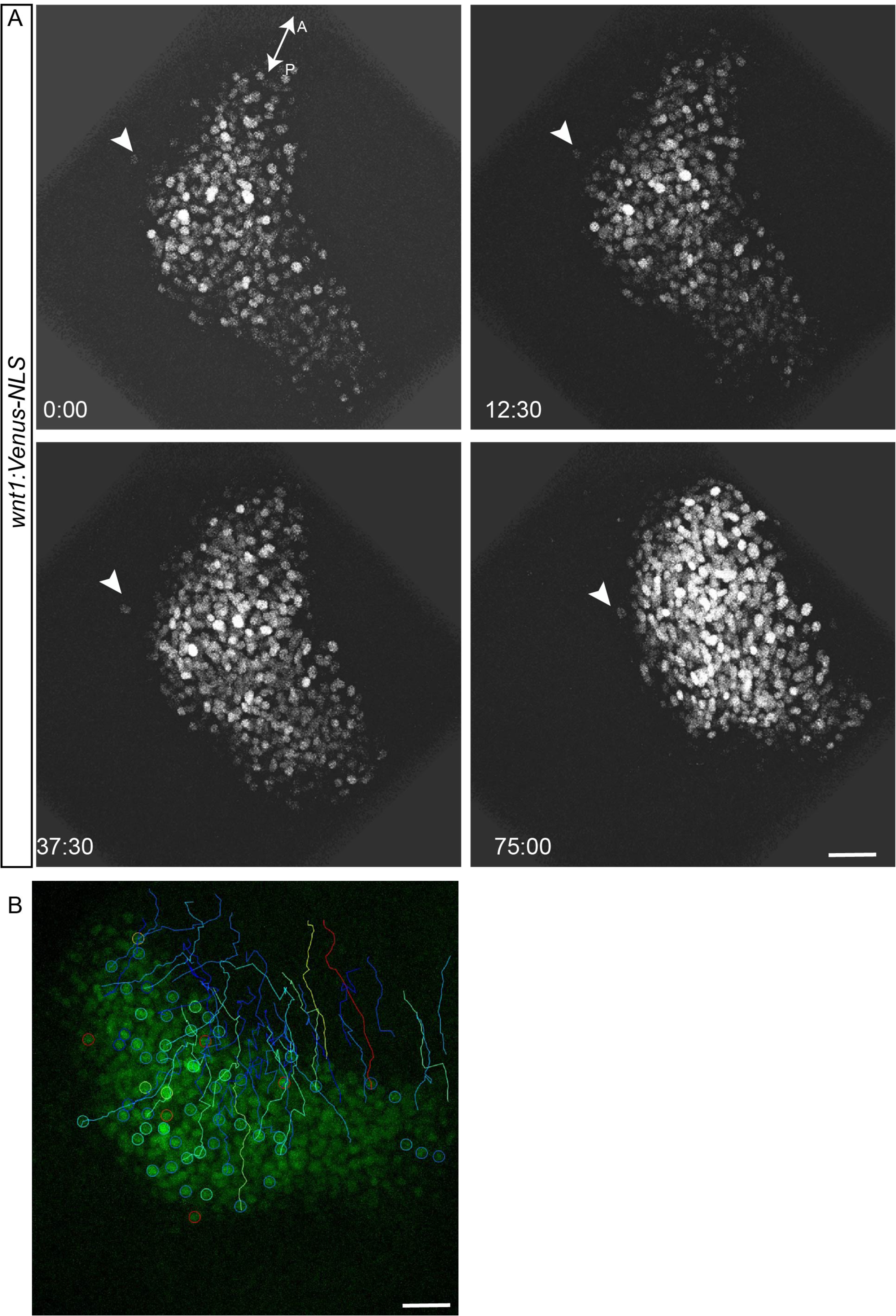
Cells sort at MHB. To visualize cell sorting during MHB development in real time, embryos from the *wnt1:Venus-NLS* reporter line were mounted and imaged dorsally between 10.5 - 12 hpf, such that the neural plate and the neural keel stages were captured. Tissue sections spanning about 30 µm were chosen with a z-interval of 1 µm. Images were acquired at 2:30 (min:sec) intervals. (A) The *wnt1:Venus-NLS*-positive cells in the midbrain are seen undergoing morphogenetic process such as neural plate convergence and migration towards the anterior end. Importantly, one *wnt1*:*Venus*-positive cell (arrowhead) that was separated from the rest of the boundary cells showed active migration towards the group of other boundary cells. Time in minutes: seconds. (B). Cell tracking showing the intermingling of cells with zig-zag movements and crossovers of tracks. Scale bar: 40 µm.

### Differential adhesion as a mechanism of cell sorting at the MHB

Proposed mechanisms of cell sorting include the differential tension hypothesis, the differential adhesion hypothesis, and the repulsion hypothesis (Batlle and Wilkinson, 2012); here, we investigated if differences in adhesion contribute to cell sorting at the MHB. Specifically, we used atomic force microscope-based single cell force spectroscopy (AFM-SCFS) (Fig 6A, schematic representation of SCFS; Krieg et al., 2008) to determine potential differences in cell-cell adhesion properties of prospective midbrain and hindbrain cells.

**Figure 6.**
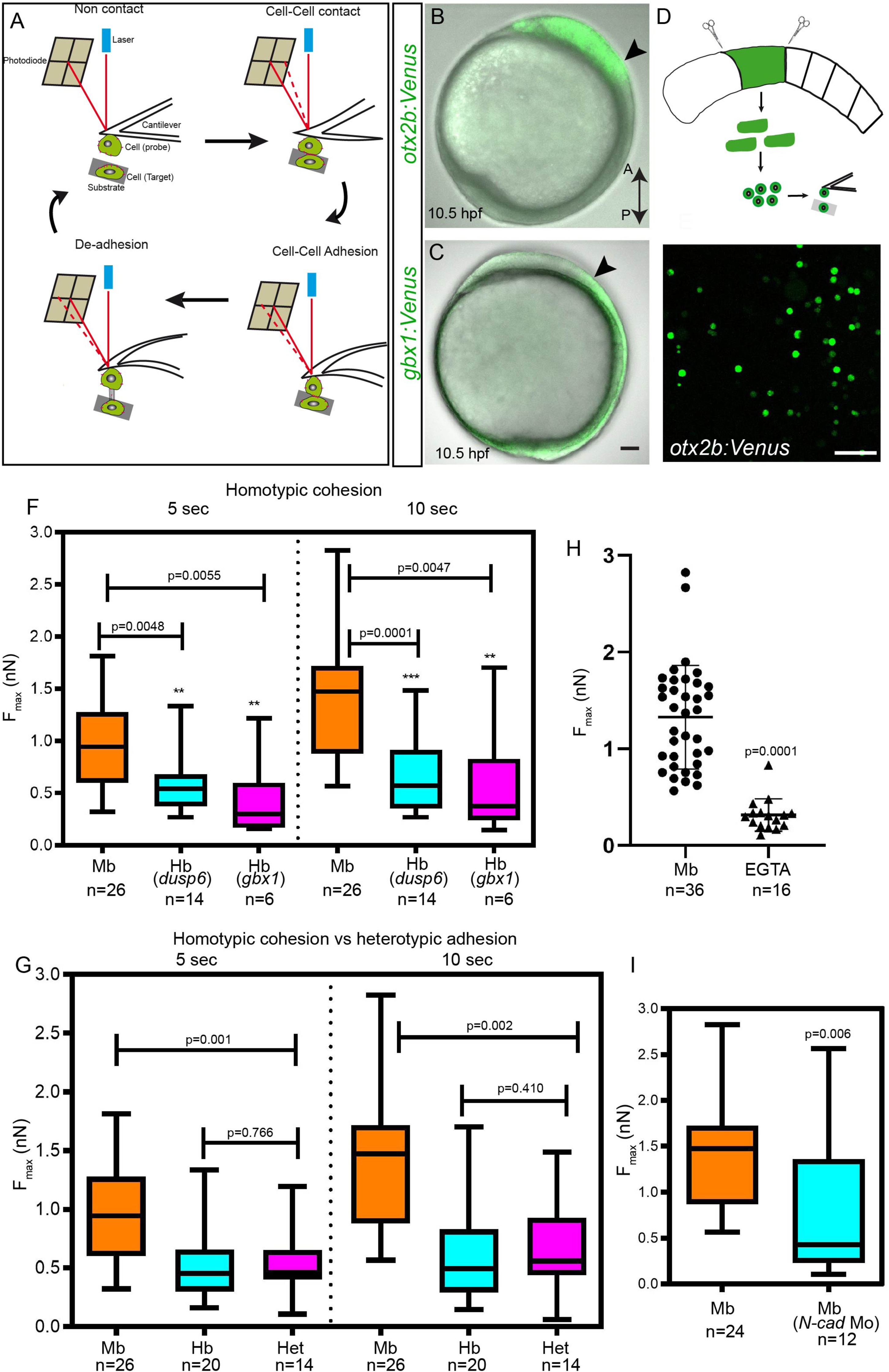
Differential adhesion as a mechanism of cell sorting at the MHB. (A) Schematic representation of atomic force microscope-based single cell force spectroscopy (AFM-SCFS). A cell attached to the cantilever is brought in contact with another cell placed on the substrate (approach). During cell-cell contact, adhesion molecules will form bonds in the contact zone. After a predefined contact time, the cantilever is retracted, which results in the breaking of bonds between the two cells (retract), and thus, the maximum force (Fmax, measured in nanonewtons, nN) required to separate the two cells can be measured (adapted from Krieg et al., 2008). (B and C) Lateral view *of otx2b:Venus* (midbrain, Mb), *gbx1:Venus* (hindbrain, hb) embryos at 10.5 hpf, arrows mark the prospective MHB. (D and E) The embryos were dissociated at the tail bud stage and a single-cell suspension for SCFS was made by mechanical trituration. (F and G) The midbrain cells (*otx2b*:Venus-positive) showed more cohesion than hindbrain cells (*gbx1* or *dusp6* positive) at the 5-second and 10-second time points (homotypic adhesion), while the heterotypic adhesion between Mb with Hb cells were lower than that of Mb-Mb cohesion. The p value (comparison between Mb and Hb; Mann-Whitney u test) for the two time points are shown on the graph; n represents the number of cell pairs analyzed for each condition. (H) Depletion of Ca2+ ions by adding 5mM EGTA dramatically reduced cell-cell cohesion between Mb cells. (I) Depletion of N-cadherin using a morpholino-mediated knockdown (N-cad Mo) reduced the cell-cell cohesion between *Otx2b* cells (midbrain cells); n represents the number of cell pairs analyzed. All the AFM based experiments were repeated at least 3 times.

Individual progenitor populations were isolated from various knock-in reporters like the *otx2b:Venus* (for midbrain cells), *gbx1:Venus* (for hindbrain cells) and *dusp6:d2eGFP* transgenic line (for hindbrain cells) (Fig 6B and 6C). Embryos were dissociated at the tail bud stage (10 hpf) and a single-cell suspension for AFM-SCFS was made by mechanical trituration (Fig 6D and 6E). Measurements of the adhesive strength of progenitors of the same kind (homotypic adhesion or cohesion) showed that the midbrain cells have greater cohesion than hindbrain cells at both contact times tested (5 and 10 seconds; Fig 6F). In contrast, adhesive forces between different cell types (heterotypic adhesion) were significantly lower than the cohesive force between two midbrain cells, but comparable to homotypic cohesion in hindbrain cells (Fig 6G). These data imply the presence of differential adhesion between midbrain and hindbrain cells.

Calcium-dependent adhesion molecules (cadherins) are known to be essential for cell-cell adhesion during embryonic development. Therefore, to determine if calcium is required for cohesion between midbrain cells, we used EGTA to chelate calcium in the media and measured cohesion using AFM-SCFS. Calcium chelation dramatically reduced cohesive strength between midbrain cells, suggesting that cohesion is calcium-dependent (Fig 6H).

Next, as N-cadherin (N-cad, *cdh2*) is the major mediator of calcium-dependent adhesion, we used morpholinos to knockdown N-cad and demonstrate that the absence of N-cad led to a signification reduction in cohesive strength (Fig 6I). Taken together, these findings suggest that the observed differential adhesion between midbrain and hindbrain cells utilizes calcium and that it is primarily mediated by N-cad in the developing neural plate.

### Molecular mechanisms of cell sorting at the MHB

To substantiate the above-described role of N-cad in cell-cell adhesion at the MHB, N-cadherin (*cdh2*) expression was perturbed by two CRISPR/Cas9-mediated knock out approaches. In the first approach, global N-cad mutants were generated by injecting 2 sgRNAs targeting exon 1 and 2, along with Cas9 mRNA, in 1-cell embryos. PCR genotyping showed efficient deletion and crispants displayed phenotypes consistent with previously described *N-cad* mutants at 24 hpf (Jiang et al., 1996, Lele et al., 2002). Importantly, *in situ* hybridization showed disorganized gene expression boundaries for *otx2b* and *egr2b* in these mutants. In contrast, control fish showed clearly demarcated boundaries for these markers at 24 hpf. Additionally, in the N-cad mutants, individual *otx2*-positive cells were visible outside their expression domain; again, such cells were absent in the control fish (Figs 7A and 7B). In the second approach, N-cad was conditionally ablated in the otx2b domain using Cre/lox-controlled Cas9 combined with the *otx2b:CreER*_T2_ driver. This conditional perturbation of N-cad in the midbrain also resulted in irregular gene expression boundaries for *otx2b*, along with the presence of *otx2b*-postive cells outside of the endogenous *otx2b* expression domain at 10 hpf (Figs 7C and D). Thus, both targeted and global N-cad deficiency yielded very similar phenotypes. Taken together, these results imply that N-cad is necessary for establishing a sharp expression boundary for Otx2.

**Figure 7.**
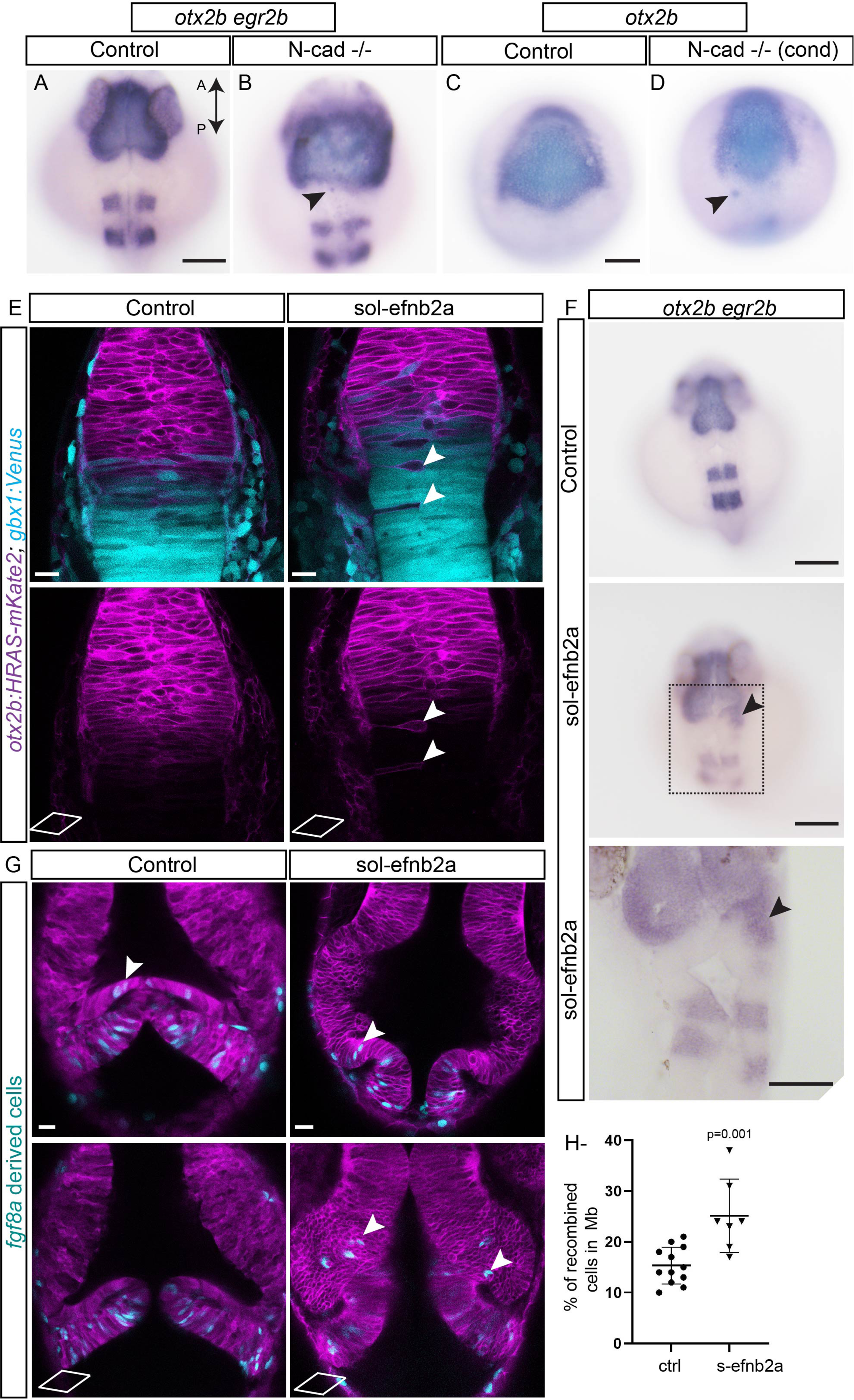
Molecular mechanisms of cell sorting at MHB. To test whether N-cadherin (*cdh2*) plays role in cell sorting, two approaches using CRISPR/Cas9-mediated mutant analysis was carried out. First, exons 1 and 2 of N-cadherin were targeted to generate global N-cad-/- embryos by injecting the sgRNA with Cas9 mRNA at the 1-cell stage. (A and B) *in situ* hybridization analysis of mutant embryos at 24 hpf for *otx2b* and *egr2b* showed disorganized pattern with fuzzy gene expression boundaries for both markers compared to control embryos. Individual *otx2b* cells outside their expression domain are seen (arrowheads). This phenotype was observed in 35% of the embryos analyzed (n=26). Second, using Cre/lox-controlled Cas9 combined with the *otx2b:CreER*_T2_ driver, N-cad was conditionally (cond) ablated in the *otx2b* domain. (C and D). Conditional perturbation of N-cad in the midbrain resulted in irregular gene expression boundaries (*otx2b*) and a few cells outside the expression domain (arrowheads). This phenotype was observed in 42% of the embryos analyzed (n=90). Soluble efnb2a (sol-efnb2a) was injected as mRNA in 1-cell stage embryos in the *otx2b:HRAS-mKate2*; *gbx1:venus* double transgenic line. (E) Perturbed Eph-ephrin signaling resulted in mis-sorting of cells across the MHB, presence of single *otx2b*-positive cells in the hindbrain (arrowheads), and *otx2b*-positive*, gbx1*-positive cells distributed further away from the Otx-Gbx overlapping domain. (F) *in situ* hybridization analysis of embryos analyzed at 24 hpf for *otx2b* and *egr2b* showed presence of *otx2b*-positive cells outside their expression domain in sol-efnb2a injected embryos (arrowheads; region marked with a dotted rectangle is enlarged in the panel below). This phenotype was observed in 44% of the embryos analyzed (n=206). (G) Embryos of the *Tg(fgf8a:CreER_T2_*); *Tg(hsp70l:loxP-DsRed-loxP-EGFPNLS)* were injected with sol-efnb2a and HRAS-mKate2 (to mark cell membrane) at the 1-cell stage. 4-OH tamoxifen mediated recombination was induced at 6 hpf. Embryos were heat shocked at 24 hpf (to label recombined cells) and imaged at 36 hpf. Perturbed Eph-ephrin signaling resulted in greater numbers of *fgf8*-derived cells in the midbrain domain (arrowheads). Two representative sections from the dorsal (top) and ventral domains (bottom) are shown. (H) Quantification of *fgf8a*-derived recombined cells in the midbrain showed an increase (ctrl vs s-efnb2a) in the s-efnb2a treated embryos. The two-tailed, unpaired ‘*t*’-test was used to calculate statistical significance, and each point in the graph represents an individual embryo (control (ctrl), n=12, s-efb2a (n=7)), p=0.0010. Scale bar: 100 µm (A-D and F), 20 µm (E and G).

The Eph/Ephrin signaling pathway is a known regulator of cell and tissue segregation during various stages of embryonic development (Xu et al., 1999). Importantly, it has been shown that the Eph receptor EphB4a is expressed in the midbrain (Cooke et al., 1997) while the ligand efnb2a is expressed in a complementary manner in the hindbrain (rhombomere1) (Cooke et al., 2005) and we have observed the same expression pattern from late gastrulation stages (8-10hpf). Therefore, to test if Eph-ephrin signaling plays a role in cell sorting at the MHB, we expressed a truncated, soluble form of Efnb2a (sol-efnb2a), which competes with various endogenous ephrin ligands that bind to Eph receptors, to perturb both forward and reverse Eph/Ephrin signaling (Cavodeassi et al., 2013; Cooke et al., 2001). Therefore, *sol-efnb2a* mRNA was injected in 1-cell embryos of the *otx2b:HRAS-mKate2*, *gbx1:Venus* double transgenic line to monitor the Otx-Gbx gene expression boundary. Perturbed Eph-ephrin signaling resulted in mis-sorting of cells across the MHB and presence of single *otx2b*-positive cells in the hindbrain. Additionally, *otx2b*-positive and *gbx1*-positive cells were seen distributed further from the Otx-Gbx overlapping domain compared to control embryos (Fig 7E). These observations were confirmed by *in situ* hybridization for *otx2b* and *egr2b*, which revealed the presence of *otx2b*-positive cells outside their expression domain in the sol-efnb2a injected embryos (Fig 7F).

Finally, we used Cre/lox-based permanent labelling to understand the effects of disrupted Eph/Ephrin signaling in lineage restriction and cell sorting at the MHB. Specifically, *fgf8a:CreER_T2_* fish were crossed with the *Tg(hsp70l:loxP-DsRed*-*loxP-EGFPNLS)* to not only visualize sorting defects, but also enumerate the number of cells that have mis-sorted. Embryos exposed to 4-hydroxy-tamoxifen at 6 hpf and imaged at 36 hpf revealed the presence of *fgf8a*-derived cells in the midbrain. Notably, when s-efnb2a mRNA was injected in these embryos at the 1-cell stage, the number of such *fgf8a*-derived cells in the midbrain was higher compared to control fish (Fig 7G and 7H). These results indicate that eph-ephrin signaling is involved in cell sorting at the MHB.

Recently, actomyosin and Yap-mediated mechanisms have also been shown to be essential for maintaining rhombomere boundaries in zebrafish (Calzolari et al., 2014.,Voltes et al., 2019). However, our analysis with F-actin reporters and active Yap signaling reporter did not reveal any involvement during MHB formation (supplementary Fig S2). Therefore, these results suggest that, at the MHB, N-cad mediates cell-cell adhesion and that Eph/Ephrin signaling is involved in cell sorting, both of which serve to establish and maintain sharp gene expression boundaries at the MHB.

## Discussion

Using multiple transgenic reporter fish for lineage tracing and live imaging, we show that lineage restriction and formation of sharp gene expression boundaries in the developing MHB occur through multiple complementary mechanisms, namely cell-fate plasticity and cell sorting, and that these processes involve differential adhesion, N-cad, and Eph-Ephrin signaling (graphical summary, Fig 8).

**Figure 8.**
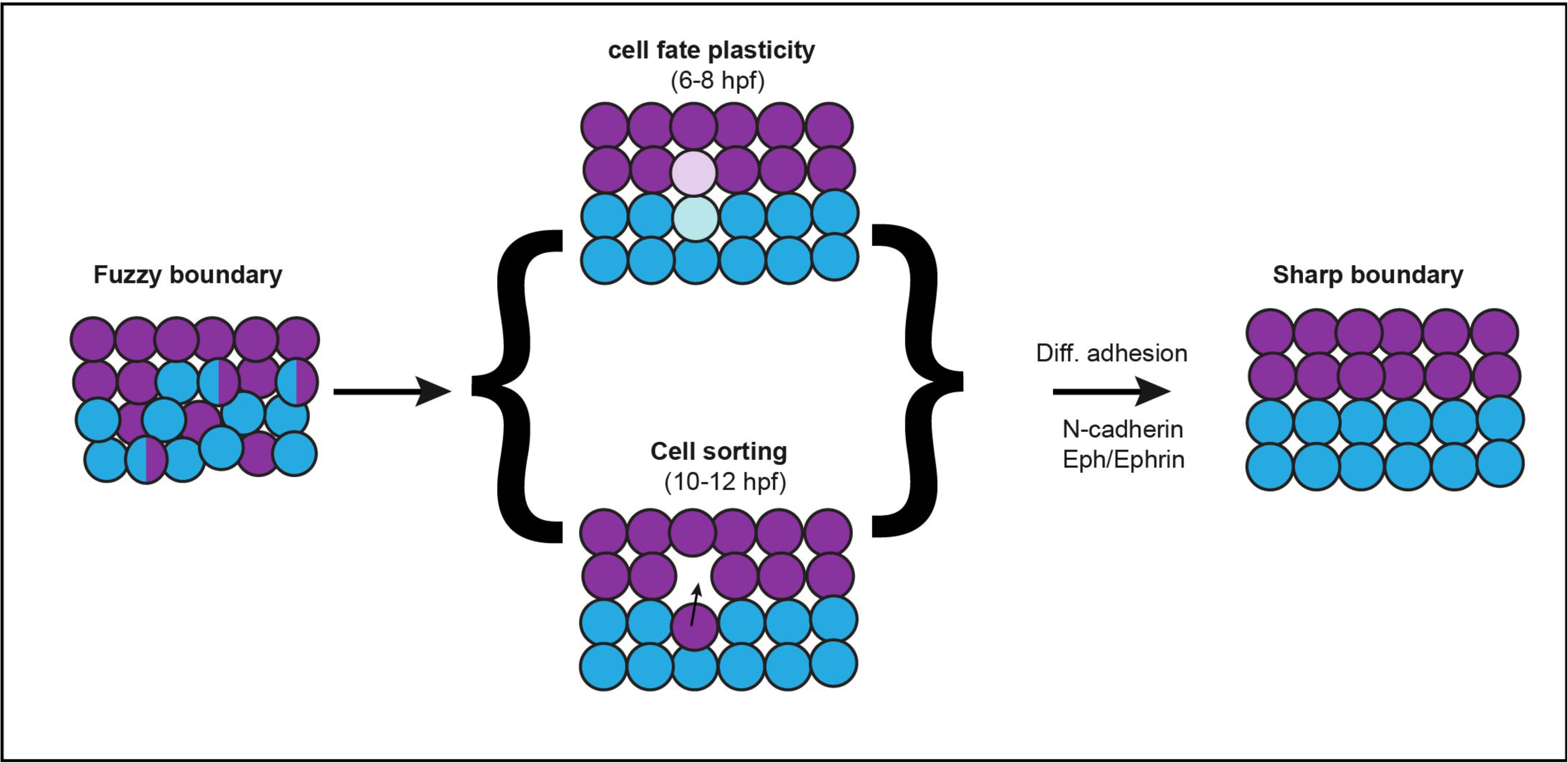
Multiple mechanisms establish sharp midbrain hindbrain boundyry. Schematic representation of how a fuzzy boundary develops into a sharp boundary over time, which involves complementary mechanisms such as cell-fate plasticity and cell sorting, involving differential adhesion, N-cadherin and Eph-Ephrin signaling.

Previously, in zebrafish, using RNA *in situ* hybridization, we found that, at 60% epiboly (6 hpf), the anterior border of the *gbx1* expression domain directly abuts the *otx2b* domain with an overlap of the two domains covering 3-4 cell layers; this structure subsequently resolves into sharply defined non-overlapping adjacent segments (Rhinn et al., 2003). Due to technical limitations (fixed samples) and the absence of reporter lines, these overlapping, and segregation events could not be visualized by live imaging, or lineage traced to follow their fate in these earlier studies. Here, using various knock-in fluorescent reporters and Cre driver lines to visualize and follow cell fate, we show that cell-fate plasticity does occur across the gene expression domains abutting the developing MHB. This phenomenon predominantly occurs during early gastrulation stages (6-8 hpf) and reduces by the end of gastrulation (10 hpf). A possible explanation for this phenomenon is as follows. Morphogens such as Wnt (in the midbrain domain), Fgf (in the hindbrain domain), transcription factors, like Engrailed1/2, and Pax2/5/8 (across the MHB), and cell adhesion molecules including Eph/Ephrin (present in adjacent domains) are expressed between 8-10 hpf. The presence of such a multitude of factors can increase cellular complexity and lead to reinforced fate commitment and lineage restriction.

Comparable results from other studies support this explanation. For example, a recent study on zebrafish hindbrain development demonstrated cell identity switching through a mechanism involving segment identity and retinoic acid signaling, with intermingling between segments and consequent cell-identity changes occurring during early stages of rhombomere segment formation i.e., before the establishment of robust Eph/Ephrin signaling that causes cell segregation across rhombomere boundaries (Addison et al., 2018). Likewise, using a combination of cell transplantation, iontophoretic cell labelling, and live-cell imaging we found that lineage restriction occurs at the MHB during the late gastrulation stages (Langenberg and Brand, 2005; Langenberg et al., 2006) in zebrafish, with similar observations being reported in chick and mice (Zervas et al., 2004, Sunmonu et al., 2011., Tossell et al., 2011). Similarly, cell lineage analysis during hindbrain development in chick has shown that the clonal progeny of cells labelled before the establishment of morphological boundaries display plasticity in cell-fate specification; in contrast, clones labelled after boundary formation are confined to their respective segments (Fraser et al., 1990).

Interestingly, we have noted a consistent pattern in our lineage tracing results, i.e., that while *gbx1*-postive cells in the midbrain domain are always also *otx2b*-postive (double positive) and are relatively fewer in number, *otx2b*-positive cells in the hindbrain domain are rarely *gbx1*-positive and relatively more in number. While there are a few potential explanations, currently, there are no known molecular mechanisms that can adequately explain this observation.

Between 10 and 12 hpf, when the neural plate transforms into the neural keel, convergent extension occurs, involving extensive cell intercalation, cell division, and intermingling of cells along the A-P axis (Kimmel et al., 1994), suggesting that mechanisms other than cell-fate switching may exist to maintain sharp expression boundaries. Consistently, using live imaging, we show that cells actively sort at the MHB between 10-12 hpf and that this sorting is influenced by N-cad-mediated adhesion between the midbrain and hindbrain cells. These findings are consistent with previous data on the role of N-cad in cell convergence and maintenance of neuronal positioning during vertebrate neural tube development is known (Lele et al., 2002).

Over the years, three major classes of cell segregation mechanisms during boundary formation have been uncovered. The first is based on differential cell-cell adhesive property (differential adhesion hypothesis) that establishes interfacial tension across boundaries (Steinberg, 2007). The second is actomyosin-based establishment of cortical tension (differential tension hypothesis) at the boundaries (Harris, 1976), and the third involves cell-cell repulsion mediated by Eph/Ephrin-like signaling molecules. These mechanisms, either individually or in combination, have been shown to establish and maintain boundaries in different tissues during development (Batlle and Wilkinson, 2012). Further, in the developing spinal cord of zebrafish, it has been recently shown that a heterogeneous population of neuronal progenitors induced by sonic hedgehog signaling sort to rearrange and form sharply bordered domains, and that this sorting mechanism is mediated via N-cadherin (Xiong et al., 2013). Here, using an AFM-based assay to measure cell adhesion properties, we show that the prospective midbrain and hindbrain cells indeed have differential adhesive strengths and that adhesion between a midbrain and a hindbrain cell is lower than that of a midbrain-midbrain cell combination. Time lapse movies of MHB formation show extensive cell movement and intermingling during which cells constantly change their partners. Intuitively, such cell behavior would require differential adhesion because this intermingling and subsequent sorting leads to compartmentalization of similar cells that ultimately form a pattern.

Additionally, our data also reveal that the cell adhesion molecule N-cadherin is involved in contributing to this adhesion. Thus, in a scenario where adhesion is disrupted due to N-cad mutations, it is expected that cells will mis-sort, and we show that conditional N-cad mutants indeed display sorting defects at the MHB. These observations highlight the importance of differential adhesion during boundary formation at the MHB.

Interactions between Eph receptor tyrosine kinase and its ligand ephrin are known to control multiple cell biological processes like polymerization of actin cytoskeleton, cadherin function, and integrin-mediated adhesion (Batlle and Wilkinson, 2012). Several Eph receptors and their ligands are expressed in complementary domains in multiple tissues during development (e.g. in the developing rhombomere boundaries), and any perturbation in the signaling leads to cell intermingling between adjacent segments (Xu et al., 1999). However, loss-of-function studies using Eph/Ephrin mutants are complicated due to redundancy among the many receptors and ligands expressed in the same tissue/cell type *in vivo* (Bush and Soriano, 2012). Nonetheless, utilizing a soluble version of the ligand efnb2a to block a wide range of Eph/Ephrin bidirectional signaling, we find that Eph/Ephrin signaling plays a significant role in establishing sharp gene expression boundaries. Further studies are required to identify the specific molecular combination of Eph receptors and ligands that mediate cell sorting at MHB.

In summary, using novel fluorescent knock-in reporters, live imaging, Cre driver-based lineage tracing, and cutting-edge cell adhesion assays and mutant analysis, we describe the process of lineage restriction at the MHB. It occurs through multiple complementary mechanisms, which are, in sequence, cell-fate specification, lineage restriction, and cell sorting; the latter involving differential adhesion, N-cad and Eph-Ephrin signaling.

## Author Contributions

G.K., and M.B conceived the project and designed experiments. G.K generated the transgenic zebrafish lines using CRISPR and generated all the experimental data, S.H. generated reagents for conditional CRISPR and Cre responder lines. G.K and M.B. wrote the manuscript.

## Acknowledgments

We are thankful to the Chen and Wente labs for providing plasmids to generate Cas9 and sgRNA mRNA (via addgene); Jens Friedrichs, Heisenberg lab, and Guck lab for help with various AFM experiments, past and present members of the Brand lab for discussions, Pablo Sanchez Quintana for help with imaging, and Dr. Vasuprada Iyengar for language and content editing. We thank Marika Fischer, Jitka Michling, and Daniela Mögel for dedicated zebrafish care. The Light Microscopy Facility, a core facility of BIOTEC/CRTD at the Technische Universität Dresden, supported this work.

## Materials and methods

### Zebrafish maintenance and breeding

Zebrafish (*Danio rerio*) embryos and adults were obtained, maintained, and raised as described previously (Brand et al., 2002; Westerfield, 2000). Embryos were staged as hours post fertilization (hpf) (Kimmel et al., 1995). The wild type strain AB was used to obtain knock-in lines, and transgenic fish lines were maintained as outcrosses. None of the larvae or adult fish showed any physiological or behavioral abnormalities.

All animal experiments were carried out in accordance with animal welfare laws of the Federal Republic of Germany (Tierschutzgesetz) that were enforced and approved by the competent local authority (Landesdirektion Sachsen; protocol numbers TVV21/2018; DD24-5131/346/11 and DD24-5131/346/12;TV T1/2019), by the institutional animal welfare committee (Tierschutzkommission der Technische Universität Dresden), and in accordance with EU directives (Directive 2010/63/ EU).

### CRISPR/Cas9-mediated knock-in lines

The knock-in lines for various genes expressed at MHB were generated and maintained as previously described (Kesavan et al., 2017; Kesavan et al., 2018). The target site sequence and primers for generating baits are provided in table 1. Transgenic animals used in this study are listed in table 2.

**Table 1:**
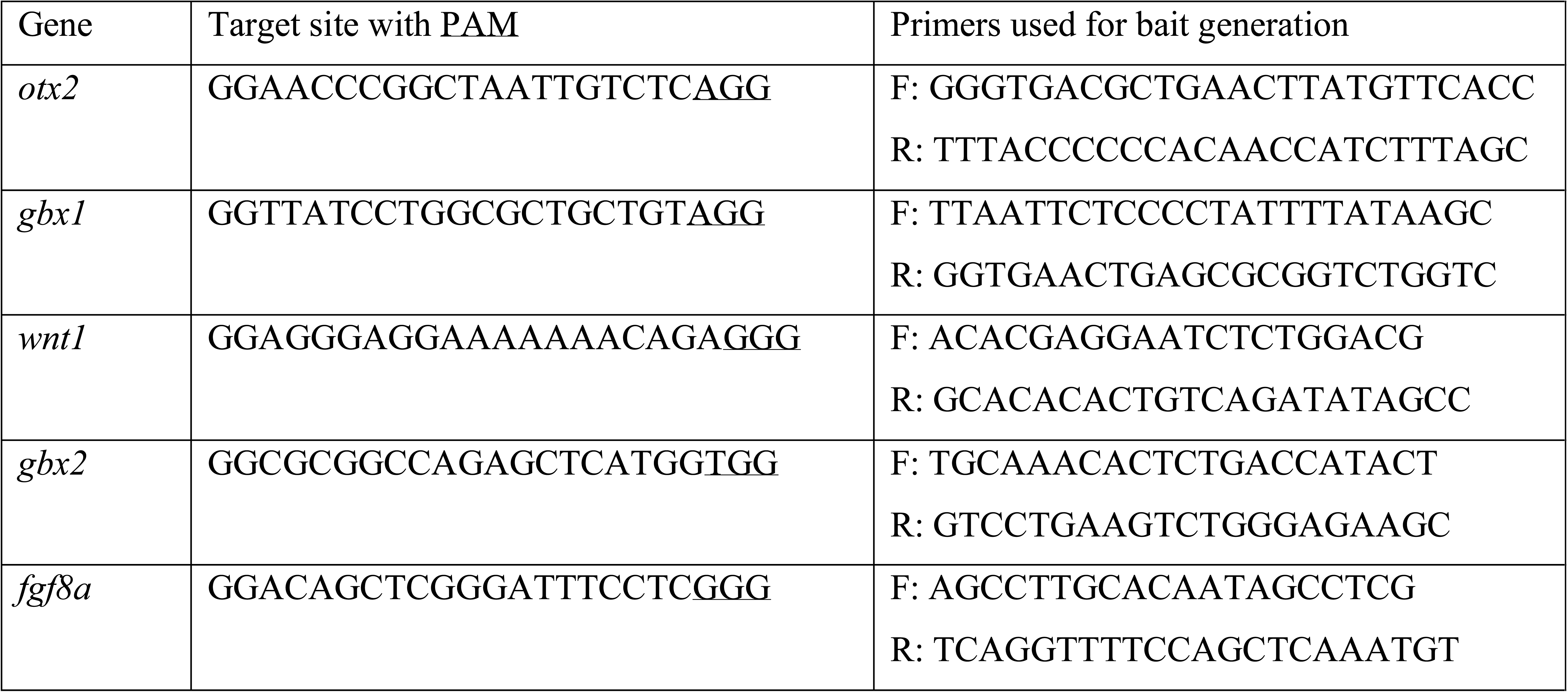
List of CRISPR/Cas9 target site and primer sequences for generating bait plasmids

**Table 2:** List of CRISPR/Cas9-mediated knock-in lines and transgenic zebrafish lines used in this study

| Strain | Source | ZFIN Identifier |
| --- | --- | --- |
| *Tg(otx2b:Venus)* | Kesavan et al., 2017 | *tud40Tg(AB)* |
| *Tg(otx2b:Venus-NLS)* | This study; Träber et al., 2019 | N/A |
| *Tg(otx2b:mKate2)* | This study | N/A |
| *Tg(gbx1:Venus)* | This study | N/A |
| *Tg(wnt1:Venus-NLS)* | This study | N/A |
| *Tg(gbx2:Venus-NLS)* | This study | N/A |
| *Tg(dusp6:d2egfp)* | Molina et al., 2007 | *Pt10Tg* |
| *Tg(otx2b:CreERT2)* | Kesavan et al., 2018 | *tud44Tg(AB)* |
| *Tg(fgf8a:CreERT2)* | This study | N/A |
| *Tg(hsp70l:loxP-DsRed-loxP-EGFPNLS)* | Knopf et al., 2011 | *tud9Tg* |
| *Tg(ubb:lox2272-loxp-RFP-lox2272-CFP-loxp-YFP)* | Pan et al., 2013 | *a131Tg* |
| *Tg(actb2:GFP-Hsa.UTRN)* | Behrndt et al., 2012 | *e116Tg* |
| *Tg(Hsa.CTGF:NLS-mCherry)* | Astone et al., 2018 | *Ia49Tg* |

### DNA, RNA, and morpholino microinjections in zebrafish embryos

mRNA for *HRAS:mKate2* or *soluble-efnb2a* were prepared using mMessage mMachine Kit (Thermofischer) and 100pg of mRNA in 1nL volume was microinjected into 1-cell stage embryos (wildtype AB). For N-cadherin morpholino experiments, 0.5pmol/embryo was injected in 1nl volume (wildtype AB). Plasmids and morpholinos used in this study are listed in table 3.

**Table 3:**
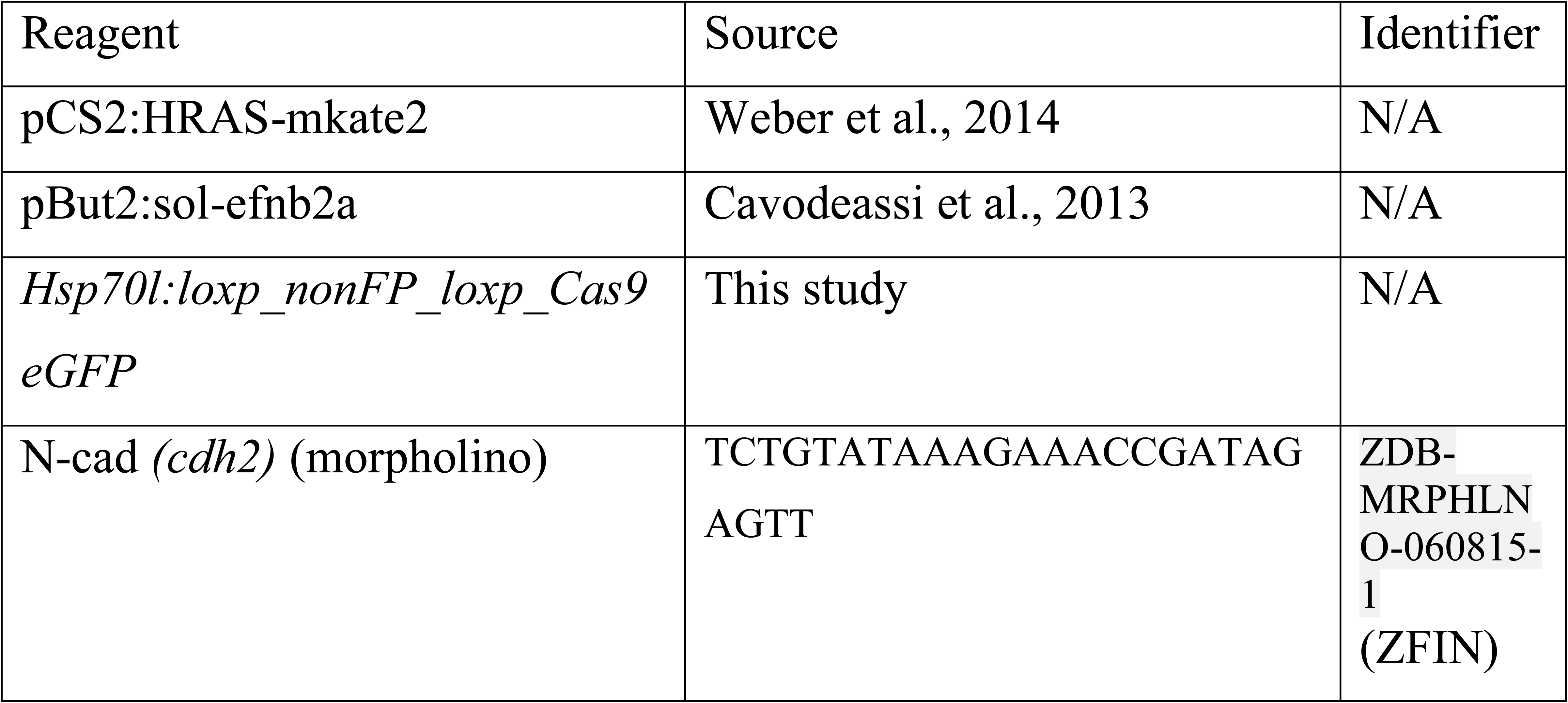
Other reagents

### Global and conditional N-cadherin mutants

For generating a global N-cadherin mutant, 1 nl of solution containing 25ng/µl of the two sgRNAs and 150 ng/µl of Cas9 mRNA was injected into 1-cell stage embryos (wildtype AB). PCR genotyping of embryos absolutely correlated with the mutant phenotype and the two sgRNA showed high efficiency in cutting the target site.

For generating the N-cadherin conditional mutants, 1 nl of solution containing 25ng/µl of the two sgRNAs and 25ng/µl of circular plasmid DNA (*Hsp70l:loxp_nonFP_loxp_Cas9eGFP*) were injected into 1-cell stage embryos obtained from *otx2:CreET_T2_*. 4-OH tamoxifen (5µM) induction was done at 4hpf and embryos were heat shocked at 8hpf for 30mins at 37°C. Embryos were sorted for eGFP fluorescence at 10.5 hpf and fixed for further analysis using *in situ* hybridization.

### *In situ* hybridization

Embryos at specific developmental stages were derived by crossing wild type fish (Ab strain). They were fixed in 4% PFA, stored in 100% methanol at −20°C, and whole mount *in situ* hybridization was performed as described elsewhere (Kesavan et al., 2017, Reifers et al., 1998). Briefly, using a RNA labeling and detection kit (Roche), digoxigenin (DIG)-labeled probes were synthesized from linear DNA, and the hybridized probes were detected using anti-digoxigenin antibody. Antibody staining was visualized using BM purple (digoxigenin). *in situ* probe staining for *otx2*, *gbx1* and *gbx2* (Rhinn et al., 2003) and *wnt1* (Lekven et al., 2003) matched patterns described previously.

### Live imaging

For live imaging, early somite stage embryos (10 to 12 hpf) were embedded in 0.7% low melting agarose. Late somite stage embryos (15 hpf onwards) were treated with 1-Pheny 2-thiourea (PTU) to block pigmentation and with MS-222 for anesthesia and mounted on a glass bottom dish (MatTek) in 1% low melting agarose. All embryos were imaged on a Zeiss LSM 780. Images were analyzed using FIJI (open source software) or Imaris (ver. 7, Bitplane), respective TIFF files generated, and figures assembled in Adobe Photoshop (ver. CS5 or CS6). For time-lapse imaging, tissue sections spanning 30-40 µm, with a Z interval of 1 µm, were imaged every 2 minutes and 30 seconds at 28°C on the LSM780 (Zeiss) microscope. The duration of time-lapse imaging was different for each imaging experiment and is mentioned in the legend of the respective figures. Maximum intensity projections of fluorescence and transmitted light images were generated using Imaris (ver. 7, Bitplane), FIJI (Schindelin et al., 2012) or Arivis 4D. Cell tracking was done using FIJI (Trackmate).

### 4-OH-tamoxifen-mediated recombination

The Cre drivers *otx2b:CreER_T2_* and *fgf8a:CreER_T2_* fish were crossed with the zebrabow responder line *Tg(ubb:lox2272-loxp-RFP-lox2272-CFP-loxp-YFP)* (Pan et al., 2013) or with *Tg(hsp70l:loxP-DsRed*-*loxP-EGFPNLS)* (Knopf et al., 2011). Embryos were treated with 1 μM 4-OH-tamoxifen (4-OHT) at 6 hpf for 12 h or with 10 μM 4-OH-tamoxifen at 24 hpf for 12 h. Control embryos were left untreated. All embryos were live imaged at 48 hpf as described above. In the zebrabow responder, a cell with multiple copies of RFP, CFP and YFP, CreER_T2_-mediated stochastic recombination events lead to cell clones being labeled by different colors. The combinatorial nature of fluorescent protein expression gives a unique barcode to each cell and marks their progenies with the same color. Untreated embryos show only default RFP expression, i.e., they represent non-recombined cells.

### Atomic force microscopy-based single cell force spectroscopy (AFM-SCFS)

The AFM-SCFS was performed as previously described (Krieg et al., 2008). AFM-SCFS is a high-precision force measuring tool that measures the difference in adhesion strength between two cells by initially bringing two isolated cells into contact. Next, after a given contact time, the cells are separated, and the force required to separate them is quantified. All experiments and data analyses used the Nanowizard I AFM setup (JPK instruments). Briefly, cantilevers (Nano world TR-TL-Au-20, nominal spring constant *k* = 32 mN m^−1^) that were coated with concanavalin A (ConA, 2.5 mg/µl, Sigma) were used. Two regions were marked in a glass-bottomed dish; one was coated with bovine serum albumin (1%) to obtain a non-adhesive substrate, while the other was coated with ConA (2.5 mg/µl) to obtain an adhesive substrate. Both substrates were gently rinsed with the HBSS (1x, life technologies) before the experiment.

Transgenic embryos staged between 10 to 10.5 hpf were carefully dissected to isolate midbrain or hindbrain progenitors, a single-cell suspension was obtained by trituration which was diluted in 1x HBSS and seeded onto the substrate. For both homotypic and heterotypic adhesion experiments, cells were identified using fluorescence microscopy, while for heterotypic adhesion experiments, fluorescent dextran, labelled with Cy5, was injected into only one set of embryos to distinguish between the two cell types. A given ‘probe’-cell was selected from the non-adhesive side of the substrate with a ConA-coated cantilever by gently pressing on it with a controlled force of 1 nN, typically for 1 s. The cell was removed from the surface for 2–10 min to allow it to firmly attach to the cantilever. The probe-cell was then moved above a ‘target’-cell that was firmly attached to the adhesive (ConA-coated) part of the substrate. Adhesion experiments (‘force-distance cycles’, see Fig. 1a) were performed at contact force of 1 nN, 10 μm s^−1^ approach and retract velocities, and contact times of either 5 or 10 seconds. Each condition (that is, same probe-target couple at the same contact time) was repeated up to three times, with a resting time of 30 s between successive contacts. Each probe-cell was used to test several target-cells and no more than 12 curves were obtained with any given probe-cell. Cells were observed continuously during and between the force-distance cycles to judge whether they were intact and were firmly attached to the cantilever or substrate. Force-distance curves were derived, and pooled data was used for statistical analysis (Graph pad prism, ver. 8). Cadherin-dependence of cell adhesion was tested after either depleting calcium by adding EGTA (5 mM, Sigma) to the medium, or by injecting embryos with morpholino oligonucleotides for *N-cadherin* (2 ng).

### Statistical Analysis

Data are presented as mean+/- SEM, unless specified otherwise. Two-tailed, unpaired ‘*t*’-test was used to calculate statistical significance at a ‘p’ value of 0.05 (Graphpad prism, ver. 8.0).

## References

1. Addison, M., Xu, Q., Cayuso, J., and Wilkinson, D.G. (2018). Cell Identity Switching Regulated by Retinoic Acid Signaling Maintains Homogeneous Segments in the Hindbrain. Developmental Cell 45, 606–620.e3.

2. Arias, A.M., and Steventon, B. (2018). On the nature and function of organizers. Development 145.

3. Astone, M., Lai, J.K.H., Dupont, S., Stainier, D.Y.R., Argenton, F., and Vettori, A. (2018). Zebrafish mutants and TEAD reporters reveal essential functions for Yap and Taz in posterior cardinal vein development. Scientific Reports 8, 1–15.

4. Batlle, E., and Wilkinson, D.G. (2012). Molecular Mechanisms of Cell Segregation and Boundary Formation in Development and Tumorigenesis. Cold Spring Harb Perspect Biol 4, a008227.

5. Brand, M., Granato, M., and Nüsslein-Volhard, C. (2002). “Keeping and raising zebrafish,” in Zebrafish: A Practical Approach, eds C. Nüsslein-Volhard and R. Dahm (Oxford: Oxford University Press), 7–37.

6. Bush, J.O., and Soriano, P., (2012). Eph/ephrin signaling: genetic, phosphoproteomic, and transcriptomic approaches. Semin Cell Dev Biol 23, 26–34.

7. Calzolari, S., Terriente, J., and Pujades, C. (2014). Cell segregation in the vertebrate hindbrain relies on actomyosin cables located at the interhombomeric boundaries. EMBO J 33, 686–701.

8. Cavodeassi, F., Ivanovitch, K., and Wilson, S.W. (2013). Eph/Ephrin signalling maintains eye field segregation from adjacent neural plate territories during forebrain morphogenesis. Development 140, 4193–4202.

9. Cooke, J., Moens, C., Roth, L., Durbin, L., Shiomi, K., Brennan, C., Kimmel, C., Wilson, S., and Holder, N. (2001). Eph signalling functions downstream of Val to regulate cell sorting and boundary formation in the caudal hindbrain. Development 128, 571–580.

10. Cooke, J.E., Xu, Q., Wilson, S.W., and Holder, N. (1997). Characterisation of five novel zebrafish Eph-related receptor tyrosine kinases suggests roles in patterning the neural plate. Dev Gene Evol 206, 515–531.

11. Cooke, J.E., Kemp, H.A., and Moens, C.B. (2005). EphA4 Is Required for Cell Adhesion and Rhombomere-Boundary Formation in the Zebrafish. Current Biology 15, 536–542

12. Dahmann, C., Oates, A.C., and Brand, M. (2011). Boundary formation and maintenance in tissue development. Nat Rev Genet 12, 43–55.

13. Dworkin, S., and Jane, S.M. (2013). Novel mechanisms that pattern and shape the midbrain-hindbrain boundary. Cell Mol Life Sci.

14. Gibbs, H.C., Chang-Gonzalez, A., Hwang, W., Yeh, A.T., and Lekven, A.C. (2017). Midbrain-Hindbrain Boundary Morphogenesis: At the Intersection of Wnt and Fgf Signaling. Front. Neuroanat. 11.

15. Harris, A.K. (1976). Is cell sorting caused by differences in the work of intercellular adhesion? A critique of the steinberg hypothesis. Journal of Theoretical Biology 61, 267–285.

16. Jiang, Y.J., Brand, M., Heisenberg, C.P., Beuchle, D., Furutani-Seiki, M., Kelsh, R.N., Warga, R.M., Granato, M., Haffter, P., Hammerschmidt, M., et al. (1996). Mutations affecting neurogenesis and brain morphology in the zebrafish, Danio rerio. Development 123, 205–216.

17. Kesavan, G., Chekuru, A., Machate, A., and Brand, M. (2017). CRISPR/Cas9 mediated zebrafish knock-in as a novel strategy to study midbrain-hindbrain boundary development. Frontiers in Neuroanatomy 11, 52.

18. Kesavan, G., Hammer, J., Hans, S., and Brand, M. Targeted knock-in of CreER_T2_ in zebrafish using CRISPR/Cas9. Cell and Tissue Research. (2018). 372(1):41–50.

19. Kiecker, C., and Lumsden, A. (2005). Compartments and their boundaries in vertebrate brain development. Nat Rev Neurosci 6, 553–564.

20. Kimmel, C.B., Warga, R.M., and Kane, D.A. (1994). Cell cycles and clonal strings during formation of the zebrafish central nervous system. Development 120, 265–276.

21. Knopf, F., Hammond, C., Chekuru, A., Kurth, T., Hans, S., Weber, C.W., Mahatma, G., Fisher, S., Brand, M., and Schulte-Merker, S. (2011). Bone regenerates via dedifferentiation of osteoblasts in the zebrafish fin. Developmental cell 20, 713–724.

22. Krieg, M., Arboleda-Estudillo, Y., Puech, P.H., Kafer, J., Graner, F., Muller, D.J., and Heisenberg, C.P. (2008). Tensile forces govern germ-layer organization in zebrafish. Nat Cell Biol 10, 429–436.

23. Landsberg, K.P., Farhadifar, R., Ranft, J., Umetsu, D., Widmann, T.J., Bittig, T., Said, A., Jülicher, F., and Dahmann, C. (2009). Increased Cell Bond Tension Governs Cell Sorting at the Drosophila Anteroposterior Compartment Boundary. Current Biology 19, 1950–1955.

24. Langenberg, T., and Brand, M. (2005). Lineage restriction maintains a stable organizer cell population at the zebrafish midbrain-hindbrain boundary. Development 132, 3209–3216.

25. Langenberg, T., Dracz, T., Oates, A.C., Heisenberg, C.P., and Brand, M. (2006). Analysis and visualization of cell movement in the developing zebrafish brain. Developmental Dynamics 235, 928–933.

26. Lekven, A.C., Buckles, G.R., Kostakis, N., and Moon, R.T. (2003). Wnt1 and wnt10b function redundantly at the zebrafish midbrain–hindbrain boundary. Developmental Biology 254, 172–187.

27. Lele, Z., Folchert, A., Concha, M., Rauch, G.-J., Geisler, R., Rosa, F., Wilson, S.W., Hammerschmidt, M., and Bally-Cuif, L. (2002). parachute/n-cadherin is required for morphogenesis and maintained integrity of the zebrafish neural tube. Development 129, 3281–3294.

28. Li, X., Zhao, X., Fang, Y., Jiang, X., Duong, T., Fan, C., Huang, C.-C., and Kain, S.R. (1998). Generation of Destabilized Green Fluorescent Protein as a Transcription Reporter. J. Biol. Chem. 273, 34970–34975.

29. Molina, G.A., Watkins, S.C., and Tsang, M. (2007). Generation of FGF reporter transgenic zebrafish and their utility in chemical screens. BMC Dev Biol 7, 62.

30. Pan, Y.A., Freundlich, T., Weissman, T.A., Schoppik, D., Wang, X.C., Zimmerman, S., Ciruna, B., Sanes, J.R., Lichtman, J.W., and Schier, A.F. (2013). Zebrabow: multispectral cell labeling for cell tracing and lineage analysis in zebrafish. Development 140, 2835–2846.

31. Reifers, F., Bohli, H., Walsh, E.C., Crossley, P.H., Stainier, D.Y., and Brand, M. (1998). Fgf8 is mutated in zebrafish acerebellar (ace) mutants and is required for maintenance of midbrain-hindbrain boundary development and somitogenesis. Development 125, 2381–2395.

32. Rhinn, M., and Brand, M. (2001). The midbrain--hindbrain boundary organizer. Curr Opin Neurobiol 11, 34–42.

33. Rhinn, M., Lun, K., Amores, A., Yan, Y.L., Postlethwait, J.H., and Brand, M. (2003). Cloning, expression and relationship of zebrafish gbx1 and gbx2 genes to Fgf signaling. Mech Dev 120, 919–936.

34. Raible, F., and Brand, M. (2004). Divide et Impera--the midbrain-hindbrain boundary and its organizer. Trends Neurosci 27, 727–734.

35. Rhinn, M., Picker, A., and Brand, M. (2006). Global and local mechanisms of forebrain and midbrain patterning. Curr Opin Neurobiol 16, 5–12.

36. Schindelin, J., Arganda-Carreras, I., Frise, E., Kaynig, V., Longair, M., Pietzsch, T., Preibisch, S., Rueden, C., Saalfeld, S., Schmid, B., et al. (2012). Fiji: an open-source platform for biological-image analysis. Nat Methods 9, 676–682.

37. Snapp, E.L. (2009). Fluorescent Proteins: A Cell Biologist’s User Guide. Trends Cell Biol 19, 649–655.

38. Steinberg, M.S. (2007). Differential adhesion in morphogenesis: a modern view. Current Opinion in Genetics & Development 17, 281–286.

39. Sunmonu, N.A., Li, K., Guo, Q., and Li, J.Y. (2011). Gbx2 and Fgf8 are sequentially required for formation of the midbrain-hindbrain compartment boundary. Development 138, 725–734

40. Tossell, K., Kiecker, C., Wizenmann, A., Lang, E., and Irving, C. (2011). Notch signalling stabilises boundary formation at the midbrain-hindbrain organiser. Development 138, 3745–3757.

41. Träber, N., Uhlmann, K., Girardo, S., Kesavan, G., Wagner, K., Friedrichs, J., Goswami, R., Bai, K., Brand, M., Werner, C., et al. (2019). Polyacrylamide Bead Sensors for in vivo Quantification of Cell-Scale Stress in Zebrafish Development. Scientific Reports 9, 1–14.

42. Voltes, A., Hevia, C.F., Engel-Pizcueta, C., Dingare, C., Calzolari, S., Terriente, J., Norden, C., Lecaudey, V., and Pujades, C. (2019). Yap/Taz-TEAD activity links mechanical cues to progenitor cell behavior during zebrafish hindbrain segmentation. Development 146.

43. Weber, I.P., Ramos, A.P., Strzyz, P.J., Leung, L.C., Young, S., and Norden, C. (2014). Mitotic Position and Morphology of Committed Precursor Cells in the Zebrafish Retina Adapt to Architectural Changes upon Tissue Maturation. Cell Reports 7, 386–397.

44. Westerfield, M. (2000) The Zebrafish Book. A Guide for the Laboratory Use of Zebrafish (Danio rerio), 4th Edition. University of Oregon Press, Eugene

45. Wurst, W., and Bally-Cuif, L. (2001). Neural plate patterning: upstream and downstream of the isthmic organizer. Nat Rev Neurosci 2, 99–108.

46. Xiong, F., Tentner, A.R., Huang, P., Gelas, A., Mosaliganti, K.R., Souhait, L., Rannou, N., Swinburne, I.A., Obholzer, N.D., Cowgill, P.D., et al. (2013). Specified Neural Progenitors Sort to Form Sharp Domains after Noisy Shh Signaling. Cell 153, 550–561.

47. Xu, Q., Mellitzer, G., Robinson, V., and Wilkinson, D.G. (1999). In vivo cell sorting in complementary segmental domains mediated by Eph receptors and ephrins. Nature 399, 267–271.

48. Zervas, M., Millet, S., Ahn, S., and Joyner, A.L. (2004). Cell behaviors and genetic lineages of the mesencephalon and rhombomere 1. Neuron 43, 345–357.

